# Experimental investigation of oxygen diffusion in the peak and valley region of minibeam patterns during X-Ray irradiation

**DOI:** 10.1101/2025.04.06.647386

**Authors:** Constantin Schorling, Evelyn Rauth, Christina Stengl, Joao Seco

**Affiliations:** Department for Physics and Astronomy, Heidelberg University, Im Neuenheimer Feld 226, Heidelberg D-69120, Germany; Division of Biomedical Physics in Radiation Oncology, German Cancer Research Center (DKFZ), Im Neuenheimer Feld 280, Heidelberg D-69120, Germany; Medical Faculty Heidelberg, Heidelberg University, Im Neuenheimer Feld 672,Heidelberg D-69120, Germany; Division of Medical Physics in Radiation Oncology, German Cancer Research Center (DKFZ), Im Neuenheimer Feld 280, Heidelberg D-69120, Germany; Heidelberg Institute for Radiation Oncology (HIRO), National Center for Radiation Research in Oncology (NCRO), Heidelberg, Germany; Radiation Oncology, Heidelberg University Hospital (UKHD), Im Neuenheimer Feld 400, Heidelberg, 69120, Germany

**Keywords:** minibeam irradiation, spatial fractionation, oxygen depletion measurements

## Abstract

**Background:** Minibeam radiotherapy has demonstrated its potential to reduce normal tissue toxicity, while maintaining tumor control. However, the underlying mechanisms behind this phenomenon remain unknown. Recent theoretical studies suggest a dose surrogate by diffusion of H_2_O_2_ into the valley regions.

**Purpose:** The aim of this study is to experimentally investigate oxygen depletion and diffusion upon minibeam (MB) irradiation.

**Methods:** A 3D-printed water phantom with four sensors was developed to enable the real-time, simultaneous measurement of oxygen concentration in the peak and valley. Water with 0 % - 11 % O_2_ and 0.1 % / 5 % CO_2_ was irradiated with broad beam (BB) and MB characterized by peak and valley widths of 2 mm x 2 mm and 0.5 mm x 2 mm. The depletion was further compared in other chemical environments.

**Results:** The oxygen depletion rates per dose in hypoxic water in the valley regions were found to be 3 to 7 times higher compared to the peaks or BB. This observation was found to be independent of oxygen concentration above 2 %, indicating oxygen depletion saturation. For MB, diffusion between peaks and valleys was observed. After a certain period, an equilibrium between diffusion and dose rate differences was established. Glutathione and HEPES as medium increased the depletion further and distinguished MB from BB.

**Conclusions:** A novel way of simultaneously measuring oxygen in the peak and valley of the MB dose pattern was introduced. The observed oxygen depletion saturation and diffusion between the peaks and valleys suggest the importance of oxygen in spatially fractionated radiotherapy studies, which is even greater for 5 mM glutathione compared to water.

## I. Introduction

In Spatially Fractionated Radiotherapy (SFRT) the irradiation is purposely blocked, creating regions of high and low doses^1–3^. Alternating high and low dose regions, also called peaks and valleys, can be achieved by introducing a collimator in the beam path ^4,5^. Distributions with peak widths in the order of 100 µm up to a few mm are called minibeams (MB)^1–3^. Recently, minibeam radiation therapy (MBRT) demonstrated a widening of the therapeutic window^1,6–11^. Animal studies revealed robust control of tumor growth in mouse models of melanoma^9^ and increased long-term survival in glioma-bearing rats^6,10,12^. Similar results were seen in glioma cells^11^. However, the rationale for tumor control in addition to normal tissue preservation with MBRT has not been sufficiently established^7,13^. One proposed mechanism is based on the production of reactive oxygen species (ROS) by water radiolysis, which disseminate from the peak into the valley regions by diffusion^14^. The production of ROS relies on the presence of molecular oxygen^15^, which is mainly expressed by two reactions^16–21^:

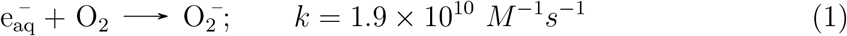

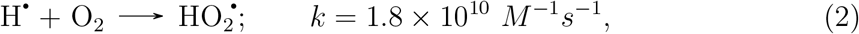

with *k* being the reaction rate constant. With 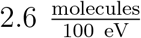 the solvate electron 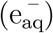 has a higher primary G-value than the hydrogen radical (H) with 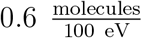. In combination with the higher reaction rate constant, 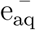 is the dominating oxygen scavenger^17,18^. The 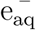 can react further with the hydroxyl radical^18–21^:

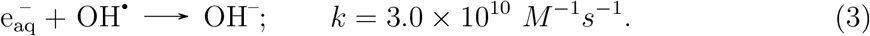

Apart from oxygen, carbon dioxide (CO_2_) is also an important parameter in physiology^22^. It is used by cells to regulate their pH, and plays a role in the respiratory system. Moreover, it has been shown that tumors in mice breathing a mixture of oxygen and CO_2_ were significantly higher perfused, increasing also the radiosensitivity^23^. CO_2_ reacts with water to form carbonic acid (H_2_CO_3_)^24–26^. However, this is only true for under 1 % of the CO_2_, while the rest stays in hydrated or aqueous form (CO_2_)_aq_^26^. The carbonic acid dissolves further into bicarbonate 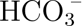 and subsequently into the carbonate ion 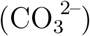.

In water saturated with 5 % CO_2_, the pH drops to 6.4^27^. Additionally, when irradiated, CO_2_ undergoes radiolysis by being split into CO and O_2_^27–29^. The by-products of the water radiolysis will react with 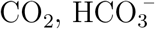 and 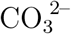^30^.

Despite the fact that chemical reactions and mechanisms have been described as surrogates for dose, the role of oxygen or carbon dioxide depletion in MB has only been the subject of theoretical investigations^14,27,31,32^. These Monte Carlo radiochemical studies were conducted in pure water; therefore, this study also uses pure water as a benchmark for other theoretical studies. As a step towards in-vitro environments, oxygen depletion was also studied in reducing and buffering agents. The aim of this work is to investigate experimentally the real-time oxygen depletion in the peaks and valleys of MB patterns for different O_2_ (0 % - 11 %, i.e. hypoxia to physioxia^33,34^), CO_2_ (0.1 % / 5 %) and MB collimators (peak x valley width: 2 mm x 2 mm, 0.5 mm x 2 mm; comparison with BB).

## II. Materials and methods

### II.A. Phantom design

The phantom was 3D printed with VeroClear (Stratasys Ltd., Israel), a PMMA like material, with inner dimensions of 30.9 mm (length) by 25.9 mm (width) by 3.91 mm (height). The transparency of VeroClear ensured optical reading with the sensors. The parts were printed on a J55 Prime 3D printer and an Objet500 Connex 1-2-3 3D printer (Stratasys, Israel). An additional sealing ring composed of Tango and VeroCyanV (Stratasys, Israel) with a Shore-A degree of hardness of 85 % was printed onto the lid to enhance the airtightness between lid and bottom part (see Fig. 1 (a)). The holders for the optical fibers were also 3D printed with VeroClear.

**Figure 1:**
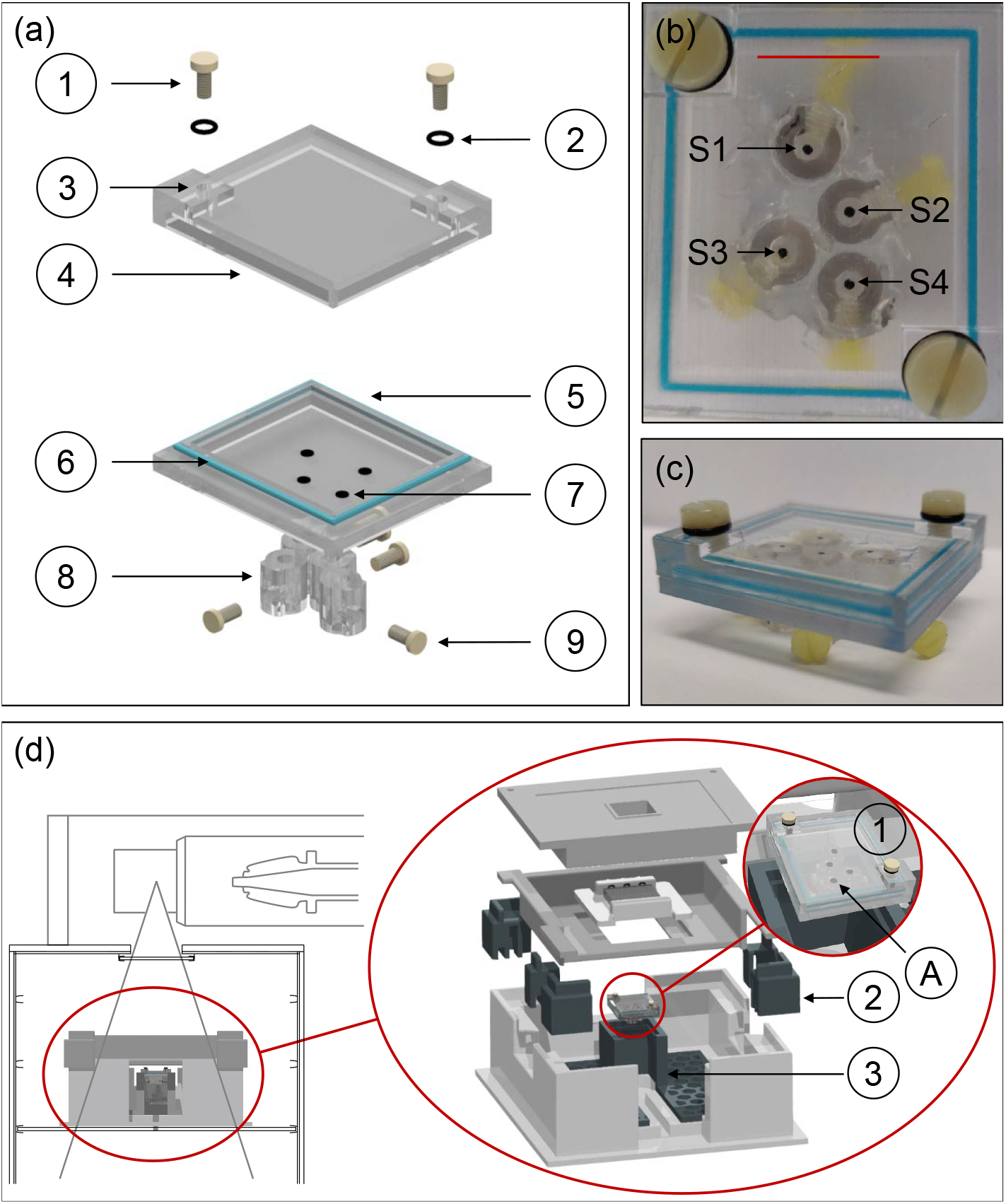
(a) Schematic of the phantom with oxygen sensors placed inside (7). M4 screws (1) and O-rings (2) close holes (3) in the lid (4), which allow for filling the phantom. The blue ring (6) around the bottom part (5) is an additional sealing layer. Holders for the optical fibers (8) are attached at the bottom. Screws (9) secure the fibers in place. (b) Top view of the phantom with the front edge (red line) of the box. The sensors 1-4 are indicated by S1-S4. (c) Side view of phantom after assembly. (d) The MB setup^2^ with the phantom (1) in the phantom holder (3) and additional adjusting components (3). During irradiation, S1 always faces the back of the setup.

### II.B. Oxygen sensors

Oxygen concentrations were measured with sensors cut from a TRACE Oxygen Sensor Foil (PyroScience GmbH, Germany) using a Cricut Joy (Cricut, USA). The mean diameter of the sensors was (1.06 *±* 0.04) mm. For readout, the optical fibers were connected to the 4-channel oxygen meter FireStingO2 (FSO2-4, PyroScience GmbH). The sensing principle was described previously by Jansen *et al*. (2021)^17^.

### II.C. Sensor placement for MB irradiation

For creating the MB dose pattern, the versatile collimator from Stengl *et al*. (2023)^2^ was used. The collimator consists of 1 mm wide tungsten plates alternating with 0.5 mm or 1.0 mm wide plastic plates to create valley and peak doses. The study investigates two different collimator configurations, with peak widths and valley widths of 2.0 mm x 2.0 mm and 0.5 mm x 2.0 mm (hereafter referred to as 2*x*2 and 0.5*x*2, respectively).

The placement of the sensors in the phantom was preceded by preliminary EBT-XD Gafchromic film measurements (Ashland, USA). Three independent measurements were performed with each collimator configuration. An in-house written script (MATLAB 2021b, MathWorks, USA), similar to Stengl *et al*. (2023)^2^, analyzed the films and calculated the mean full width at half maximum (FWHM) and center-to-center distance (CTC). Consequently, the sensors were positioned to allow measurement of both collimator configurations with a single phantom, with two sensors associated with the peak and valley regions respectively. The sensor positions were verified (see supplementary S.II. Fig. S1), which revealed a good alignment for all sensors except sensor S3 in the 2*x*2 pattern, which was therefore excluded. The sensor positions (1-4) were determined to be 9.28 mm; 15.37 mm; 19.44 mm and 22.40 mm to the front edge of the phantom (see Fig. 1 (b), 1 (c)). The uncertainty was approximated to be below 0.15 mm.

### II.D. Irradiation setup

The irradiation was performed at a Faxitron MultiRad 225 X-ray irradiation system (Precision X-Ray, USA). A phantom holder was introduced to the setup from Stengl *et al*. (2023)^2^ as displayed in Fig. 1 (d). The collimator was placed at a source-to-collimator distance of 21.8 cm, while the source-to-phantom distance was 23.9 cm. In order to ensure the correct position of the sensors in the peak and valley, a film was placed under the phantom during each MB irradiation. The phantom was irradiated with 200 kV tube voltage and 17.8 mA current. The dosimetry for the setup was performed similar to Stengl *et al*. (2023)^2^ (see supplementary section S.I.). Furthermore, sensor-specific dose rates were obtained. The phantom mean dose rate was obtained by integrating the dose over the whole phantom size and dividing the dose integral by the size. Comprehensive information concerning the sensor placement and sensor-specific dose rates can be found in the supplementary section S.II.

### II.E. Measuring oxygen depletion during irradiation

Double deionized water (Barnstead GenPure, Thermo Scientific, Germany) was placed in a Sci-Tive hypoxic chamber (Baker Ruskinn, Ruskinn Technologies Ltd., UK), where nitrogen serves as air substitute, for at least two days to achieve (1.0 - 11.0) % O_2_ and (0.1 / 5.0) % CO_2_. To mimic a radical scavenging and reducing intracellular environment, the oxygen depletion in different chemicals were investigated: Glutathione (GSH) (Sigma-Aldrich, Germany) was chosen to be investigated, as it is one of the primary intracellular scavengers and reducing agents^35^. The concentration was set to be 5 mM in accordance to usual appearance in the human body^36^. It was further rectified to a pH level of approx. 7. Additionally, the oxygen depletion was measured in HEPES (2-[4-(2-hydroxyethyl)piperazine-1-yl]ethanesulfonic acid, 10 mM) (Sigma-Aldrich, Germany) to represent buffering agents. Both reagents were brought into the hypoxic chamber at 1 % O_2_ and 5 % CO_2_. An overview of the O_2_ and CO_2_ settings for the different measurements can be found in Tab. 1.

**Table 1:** Summary of the measurement settings. The conenctration of HEPES as medium was set to 10 mM and for GSH it was 5 mM.

| Figure | Chemical | O<sub>2</sub> [%] | CO<sub>2</sub> [%] | Collimator |
| --- | --- | --- | --- | --- |
| Fig. 2 | H<sub>2</sub>O | 1 - 14 | 0.1 | — |
| Fig. 3 | H<sub>2</sub>O | 1 | 0.1 | BB, 2x2, 0.5x2 |
| Fig. 4 | H<sub>2</sub>O, HEPES, GSH | 1 | 5.0 | BB, 2x2, 0.5x2 |
| Fig. 5 (a) | H<sub>2</sub>O | 0 - 10 | 0.1, 5.0 | BB |
| Fig. 5 (b) | H<sub>2</sub>O | 0 - 5 | 0.1 | BB, 2x2 |
| Fig. 5 (c) | H<sub>2</sub>O | 0 - 5 | 0.1 | BB, 0.5x2 |

A two-point calibration of the sensors and cables was performed in normoxic and hypoxic water prior to the measurements. The raw values were subsequently corrected for temperature and pressure. After filling the phantom, a rest period was introduced to ensure stable conditions and to establish a steady state prior to irradiation. The phantom was then irradiated with BB, 2*x*2 and 0.5*x*2 MB. The oxygen depletion was measured during irradiation at a sampling rate of approximately one measurement per second. All measurements performed at approximately 1 % O_2_ were repeated three times, whereas the oxygen measurements for different initial oxygen concentrations were performed only once. Under hypoxic conditions, the phantom was irradiated for 20 min, while for varying oxygen concentrations, only 5 min BB and 2*x*2 MB intervals and 15 min intervals for 0.5*x*2 were assigned to an oxygen start value. The oxygen depletion gradients were obtained by fitting a linear function to the mean curves, subsequent to the attainment of equilibrium. The gradients were corrected for background variations that were recorded 120 s prior to each irradiation.

## III. Results

### III.A. Dose rates at the sensor locations

The BB dose rate was established to be (6.12 *±* 0.04) Gy/min. Furthermore, the dose rates for each sensor position and collimator pattern were determined (see Tab. 2). The (sensor-associated) peak-to-valley dose ratio (PVDR) for 2*x*2 is approximately five, while for the 0.5*x*2 collimator it is close to 10. In addition, the average phantom dose rate for 0.5*x*2 is less than 0.5 Gy/min, while for 2*x*2 it is three times higher at approximately 1.5 Gy/min.

**Table 2:** Summary of the sensor dose rates for the different collimator configurations. Doses were measured with EBT-XD films and plotted against irradiation time to obtain the dose rates. S3 in 2*x*2 was excluded because it was not perfectly allocated in either a peak or a valley.

| Collimator Configuration | Sensor | Dose rate [Gy/min] |
| --- | --- | --- |
| BB | S1-S4 | $6.120 \pm 0.040$ |
| 2.0 mm x 2.0 mm | S1<sup>2x2</sup> (Peak) | $3.158 \pm 0.032$ |
| | S2<sup>2x2</sup> (Valley) | $0.571 \pm 0.041$ |
| | S4<sup>2x2</sup> (Peak) | $2.838 \pm 0.060$ |
| | Mean (whole phantom) | $1.517 \pm 0.015$ |
| 0.5 mm x 2.0 mm | S1<sup>0.5x2</sup> (Valley) | $0.110 \pm 0.011$ |
| | S2<sup>0.5x2</sup> (Valley) | $0.128 \pm 0.012$ |
| | S3<sup>0.5x2</sup> (Peak) | $1.507 \pm 0.022$ |
| | S4<sup>0.5x2</sup> (Peak) | $1.281 \pm 0.035$ |
| | Mean (whole phantom) | $0.457 \pm 0.008$ |

### III.B. Oxygen depletion with BB and MB irradiation

Stability measurements without irradiation showed no random diffusion, sensor drift or high background noise throughout the measurement period for all O_2_ concentrations examined (Fig. 2). Additionally, the stability was determined in water, HEPES and GSH at CO_2_ and showed a similar conditions (not shown).

**Figure 2:**
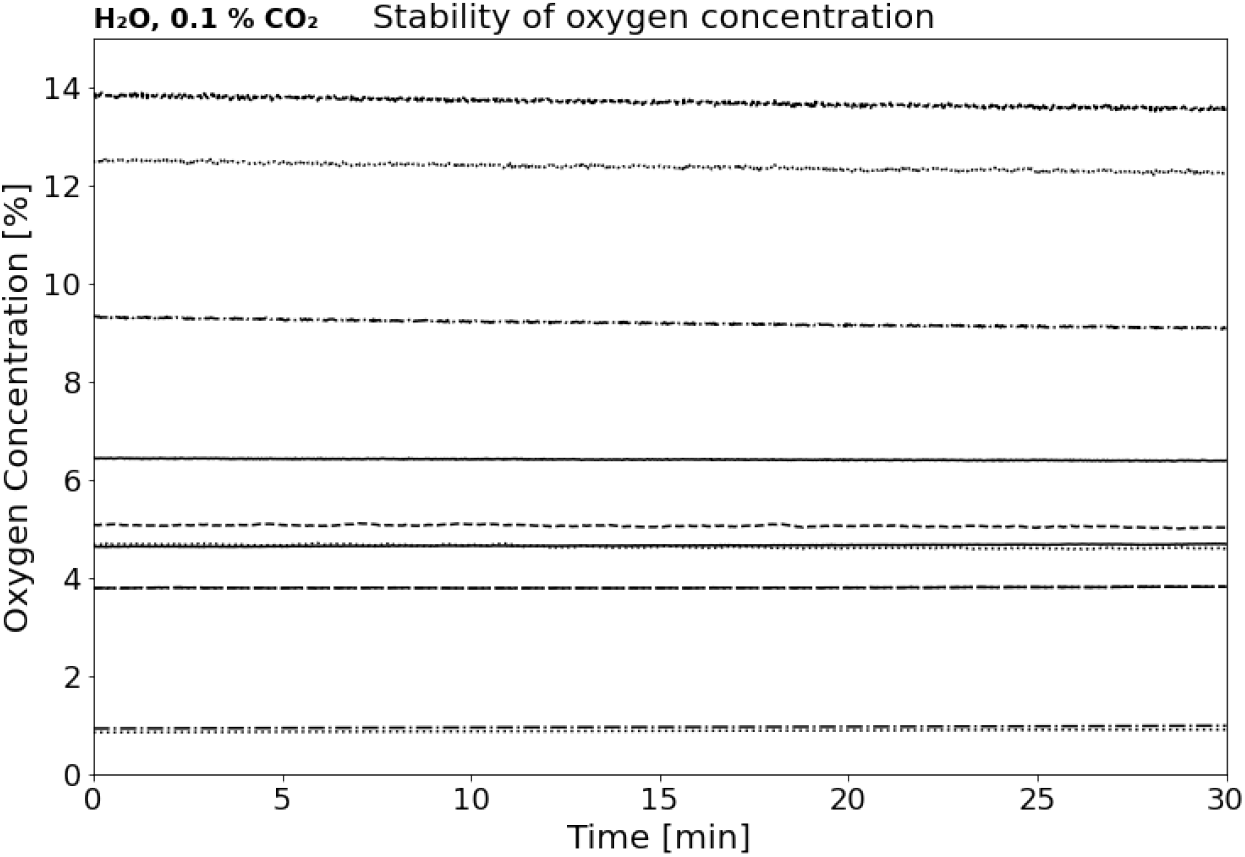
Stability measurements without irradiation of the phantom. For each background measurement (i.e. each line), the water was brought to the specific O_2_ concentrations using the hypoxic chamber.

Firstly, the oxygen depletion was studied in dependence on the spatial beam structure in hypoxic water (1 % O_2_) devoid of carbon dioxide (0.1 % CO_2_). For better comparison, all subsequent hypoxic oxygen depletion measurements were initially shifted to start at exactly 1 % O_2_. This is a reasonable simplification as all chemicals were initially brought to 1 % O_2_ and observed deviations were small. The displayed results in Fig. 3 depict each the mean and standard deviation of three independent measurements. For BB irradiation, all sensors were further averaged. The oxygen depletion as a function of irradiation time (Fig. 3 (a), 3 (c)) can be converted into a dose dependent value by multiplying time with the specific sensor dose rates (Fig. 3 (b), 3 (d)). While the peak and BB irradiation cause a sharp drop in oxygen concentration, the valley regions react in a curved manner. Subsequent to an initial equilibration phase, oxygen depletion is found to be mostly linear for all beam structures and sensor locations. Following this, a linear fit was applied after equilibration to obtain the oxygen depletion gradient. Due to the high dose rate, the BB has a high oxygen depletion over time, but not per dose. Although the 2*x*2 peak dose rates for the MB patterns are much lower than for the BB, their oxygen depletion per dose is similar to that of the BB. For the 0.5*x*2 peak sensors it is half. Compared to BB, the oxygen depletion per time for the valley sensors is low, however, they receive even less dose, resulting in oxygen depletion rates per dose that are 3 to 7 times that of BB.

**Figure 3:**
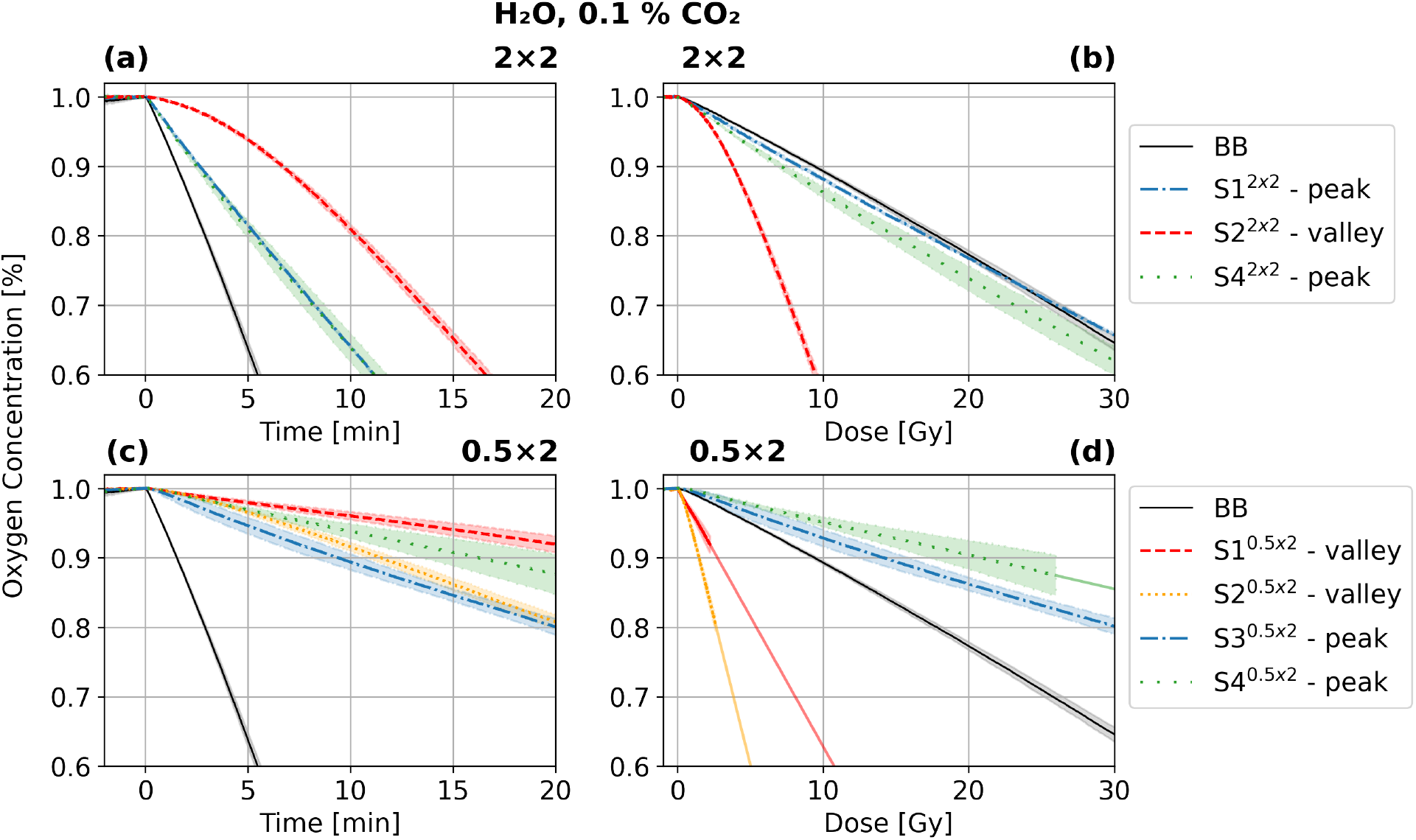
Mean oxygen depletion in hypoxic water (1 % O_2_, 0.1 % CO_2_) for each sensor over time (a), (c) and dose (b), (d) for BB, 2*x*2 and 0.5*x*2 irradiation. The phantom was irradiated for 20 min. For S1^0.5*x*2^, S2^0.5*x*2^ and S4^0.5*x*2^ the dose dependent oxygen concentration was extrapolated by a linear line based on the depletion from the applied last 1 Gy irradiation.

To compare the results obtained for water with different environments, the oxygen depletion per dose was measured and calculated in the same way as for the hypoxic water with 0 % CO_2_ (Fig. 4 (a)-4 (f)). It is evident that the gradients observed are comparable to those of hypoxic water without CO_2_. For water with 5 % CO_2_ the BB depletion was not significantly altered, in the 2*x*2 pattern only S1 (peak) showed a 6 % slower depletion. For 0.5*x*2 the oxygen depletion of the peak sensors was increased, while for the valley sensors it was reduced, but this trend is only significant for S3 (peak).

**Figure 4:**
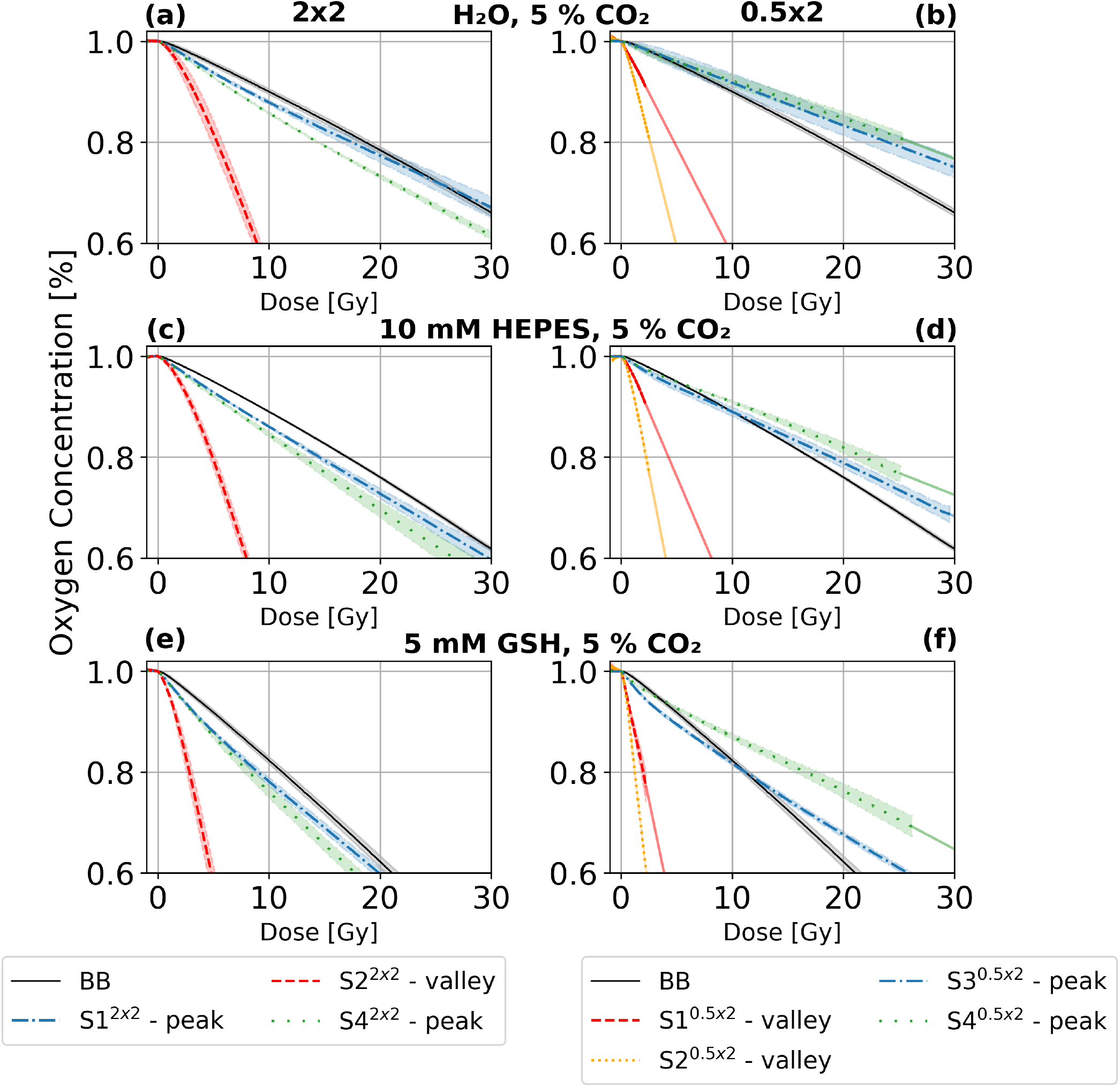
Mean oxygen concentration dependent on applied dose for different environments: H_2_O at 1.2 % O_2_ and 5 % CO_2_ in (a) and (b); HEPES at 0.6 % O_2_ and 5 % CO_2_ in (c) and (d); GSH at 0.8 % O_2_ and 5 % CO_2_ in (e) and (f). The phantom was irradiated for 20 min. For S1^0.5*x*2^, S2^0.5*x*2^ and S4^0.5*x*2^ the dose dependent oxygen concentration was extrapolated by a linear line based on the depletion from the applied last 1 Gy irradiation.

For HEPES at 5 % CO_2_, oxygen depletion was significantly increased for all sensors and irradiation structures. While the relative deviation for the BB irradiation is still small at about 4 %, it becomes more significant at 12 % to 57 % for the MB irradiation. In general, a higher depletion amplification was observed for the peak sensors than for the valley sensors, so that the peak-valley deviation gap was reduced. For GSH at 5 % CO_2_, a significant increase in depletion was found, with at least 57 % higher depletion rates. Here the valley sensors detected an even higher increased depletion than the peak sensors.

The oxygen depletion per dose can be plotted against varying oxygen concentrations and fitted with Michaelis-Menten kinetics (Fig. 5 (a)-5 (c))^37^. In this context, the substrate concentration is associated with *c*(*O*_2_), and *V*_*max*_ with the maximum rate of oxygen depletion, resulting in 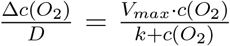. For BB and oxygen concentrations above 2 %, *c*(*O*_2_)*/D* is found to be independent of the oxygen concentration, regardless the CO_2_ concentration. Furthermore, at 5 % CO_2_, there is a 7 % higher depletion for physioxic conditions. Similar to BB, both, peak and valley of the MB exhibit a plateau for *c*(*O*_2_)*/D* above 2 % O_2_. In contrast, the valleys demonstrated a significantly higher oxygen depletion per dose. This phenomenon is particularly evident in the 0.5*x*2 valleys. *c*(*O*_2_)*/D* of these valleys are 4 to 9 times higher than those of the peaks and the BB, for 2*x*2 it is almost 5 times higher.

**Figure 5:**
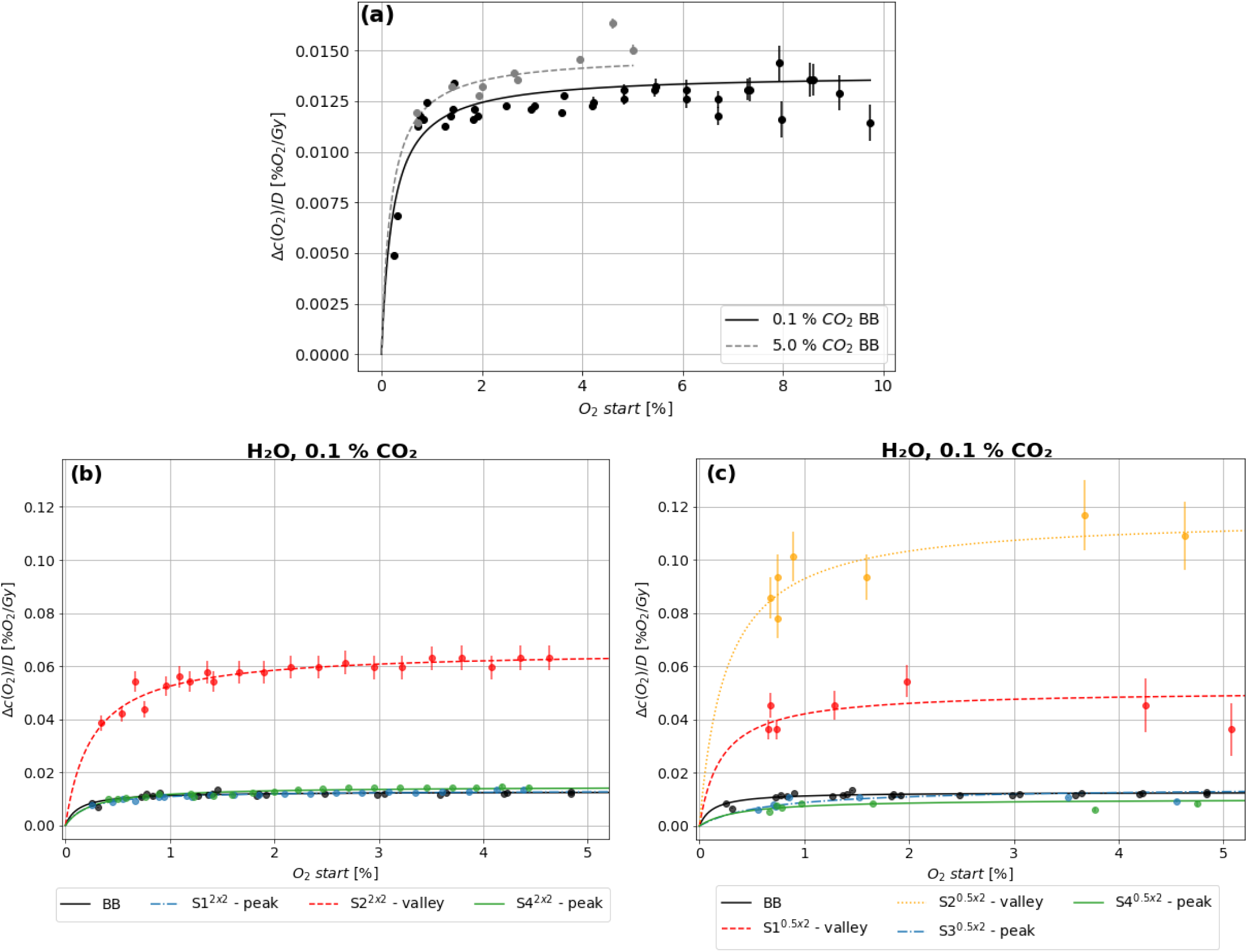
Michaelis-Menton fits of oxygen depletion per dose as a function of O_2_ start. The Michaelis-Menton fits for water with 0 % CO_2_ and 5 % CO_2_ irradiated with BB were compared (a); in addition, for 0.1 % CO_2_ BB was compared with 2*x*2 (b) and 0.5*x*2 (c) collimator patterns.

## IV. Discussion

This study focused on the simultaneous measurement of oxygen concentration in the peaks and valleys of a MB pattern during irradiation. Although the effect of MB irradiation on animals has already been subject to studies ^5,10,38^ and there are several hypotheses, the underlying mechanism of the minibeam effect is still unknown^1,8,14,39,40^. The minibeam effect has been shown for collimator configurations with the valley width being two to four times larger than the peak width^2,5,7,41,42^. Therefore, the 0.5 mm x 2.0 mm and 2.0 mm x 2.0 mm collimator configurations were chosen. The positioning of the sensors in the peaks and valleys was successful, albeit with the exclusion of S3. This is corroborated by (sensor-associated) PVDR of 5 to 14, which are consistent with the findings reported in the relevant literature^2,42^. The observed stability of the oxygen concentration in the absence of irradiation is consistent with the findings reported in the study by Jansen *et al*.^17^, in which a phantom fabricated from the same material was used.

### IV.A. Oxygen depletion in hypoxic water

In the analysis of BB irradiation, a linear oxygen depletion gradient was observed from the onset. This observation aligns with the reported findings^13,43,44^. These studies indicated a linear relationship between the duration of irradiation and oxygen depletion, irrespective of the specific irradiation type. Furthermore, the sensors exhibited a uniform response, which was anticipated given the homogeneous irradiation conditions. In the 2*x*2 configuration, there was a pronounced quadratic trend in the initial minutes for the peak sensors, and a particularly distinct inverse quadratic trend for the valley sensors. Similar observations can be seen for the 0.5*x*2 pattern. This phenomenon cannot be attributed solely to closed, non-interacting peak and valley systems, characterized by a constant dose rate difference, as these systems would also be expected to show a linear oxygen depletion, as shown for BB throughout the whole measurement period. Subsequently, after approximately seven minutes, a linear trend emerged, suggesting that oxygen depletion gradients resulting from dose rate differences are compensated by diffusion. The 2*x*2 valley is characterized by about 5 times higher oxygen depletion per dose than the peaks, and the PVDR is also approximately fivefold. For 0.5*x*2 the valley gradients are 6 to 12 times higher than the peak ones, whereas the PVDR is 10 to 13.5 times higher. Conclusive deductions concerning the diffusion process necessitate a comprehensive understanding of dose-rate-specific oxygen depletion, a concept that was previously explored by Jansen *et al*.^17^. However, the study’s primary focus was on ultra-high dose rates, with the lowest investigated dose rate being approximately 0.1 Gy/s, which aligns with our BB irradiation. Nevertheless, the MB dose rates examined in our study range from 1/2 to 1/55 of this. Indeed, that the gradients of the 2*x*2 peaks are of the order of BB and the 0.5*x*2 peaks are lower is considered to be the result of dose rate dependencies as well as diffusion, which is dependent on PVDR and peak and valley width. The disparities observed among the peak sensors and between the valley sensors are associated with the position on the phantom (see supplementary section S.II. Fig. S1). The oxygen concentration per dose for the valleys of 0.5*x*2 is extrapolated in Fig. 3 and Fig. 4 as at this positions 0.4 % O_2_ were not depleted within 20 min, which demonstrates again the low dose rates in the valleys.

### IV.B. Oxygen depletion in other chemical environments

In the presence of CO_2_, the pH value of the water changes and it becomes acidic^27^. An acidic environment favors H^•^ which is a scavenger for OH^•27^. According to reaction 3, there will be more 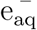 present when there is less of the hydroxyl radical. Subsequently, the higher amount of available 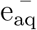 leads to an increase in oxygen scavenging according to reaction 1, and therefore a faster decrease in oxygen. Furthermore, CO_2_ and its by-products from radiolysis and dissolving into carbonic acid can also scavenge OH^•^ (see supplementary section S.III.), which adds to the described chain reaction leading to a faster decrease in oxygen. However, one of the radiolysis products of CO_2_ is oxygen^27^, increasing the amount of oxygen again and thus slowing the oxygen depletion down. In addition, CO_2_ and its by-products from radiolysis and dissolution into carbonic acid can react with 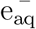^18,30,45^ (see supplementary section S.III.), which leaves less 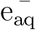 for the reaction with oxygen according to reaction 1. Therefore, the decrease in oxygen would be slowed down. We found non-significant changes in the oxygen depletion per dose of hypoxic water with 5 % CO_2_ irradiated with BB, compared to water without CO_2_ (Fig. 4 (a)-4 (b)). It is noticeable that for 0.5*x*2 the peaks tend to deplete faster, while the valleys show the opposite behavior. The increased depletion in the peaks is thought to be caused by increased 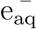 availability and additional OH^•^ scavenging. In contrast, in the valleys, oxygen depletion is reduced at 5 % CO_2_ because CO_2_ and its by-products react with 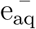, limiting its availability for oxygen reduction. However, it should be noted that no significant changes were found for 2*x*2, where peak and valley widths are suspected to be key contributors and need to be investigated further.

As a step towards in-vitro studies, oxygen depletion was investigated in a radical scavenging and reducing intracellular environment (GSH) and a buffering agent (HEPES). While the oxygen depletion for BB in HEPES was the same at 5 % CO_2_ (Fig. 4 (c)-4 (d)) compared to water without CO_2_, this changed for MB. Here the sensors detected an increase of between 12 % and almost 60 %, with the peaks tending to exceed the valleys. HEPES competes with oxygen for 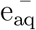, but CO_2_ can also scavenge OH^•^, leading to a complex interplay of reactions. As ROS production is generally less pronounced in the valley regions, and HEPES is less reactive to other radicals than 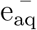, smaller increases may be seen while the peaks are more pronounced.

As GSH is a reducing agent, it is expected to be oxidized to GSSG while reducing H_2_O_2_ and scavenging OH^•^, H^•^ and 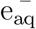 ^46–48^. We observed an increase in oxygen depletion per dose for both BB and MB, as expected^48^(Fig. 4 (e)-4 (f)). Interestingly, while oxygen depletion was increased by approximately 60 % for BB and most peak sensors, it was increased even further in all MB valleys, suggesting that the presence of GSH enhances oxygen depletion more efficiently in low-dose regions, likely due to its interaction with radiolytic species such as 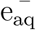 and OH^•^. This could indicate a stronger radical-driven oxygen depletion effect in valleys, where oxidative stress is more influenced by secondary reactions rather than direct ionization events.

### IV.C. Oxygen depletion in water at varying concentrations

Compared to hypoxic water, Δ*c*(*O*_2_)*/Gy* increases for oxygen concentrations up to 2 % and reaches then a plateau for physioxic and normoxic conditions (Fig. 5 (a)-5(c)). This is independent of the spatial beam structure and position within it. Jansen *et al*.^37^ found similar results for ultra high dose electron BB irradiation. The magnitude of the gradients for the BB can further be verified, as another study by Jansen *et al*.^17^ described similar gradients in this dose rate range. At a CO_2_ content of 5 %, a 7 % higher oxygen depletion per dose was found with BB for water with c(*O*_2_) above 2 %. Under these conditions, this suggests a preference for the reaction 1 over O_2_ production by CO_2_ and 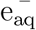 scavenging. The depletion per dose for the valleys follows that of BB and peaks at a high level. Again, no oxygen concentration dependence was found above 2 %. The deviations of the two valleys in 0.5*x*2 correspond to the position of the sensors on the phantom: S1 is located furthest outwards and is surrounded by smaller peaks than S2 (see also supplementary section II.).

### IV.D. Limitations

A limitation of this study is the placement of the sensors directly in the peaks and valleys of the patterns. Although this was done successfully and clear differences were found, with smaller sensors and even more precise placement in the peaks and valleys, even greater differences between them can be expected. While the behavior of buffering and reducing agents compared to water was examined, these environments do not fully capture the complexity of a cell and cell conglomerate. This is particularly true for oxygen supply and diffusion dynamics with biological, non-irradiated adjacent environments. This will require further research to confirm or disprove whether the radiochemical results obtained can be applied to biological systems.

## V. Conclusion

This is the first study to simultaneously measure oxygen depletion in the peak and valley of a MB pattern during irradiation. Two collimator configurations with peak and valley widths of 2.0 mm x 2.0 mm and 0.5 mm x 2.0 mm were investigated. The oxygen depletion rates per dose in the valley regions were found to be significantly higher than in the peaks and with BB. This observation was predominantly independent of oxygen concentration and chemical environment. Furthermore, the occurrence of a quadratic dose response suggests a significant role for diffusion in the MB pattern as a whole, which subsequently reaches equilibrium with dose rate differences between the peaks and valleys. The novel findings on oxygen depletion saturation and diffusion between the peak and valley regions indicate an importance of O_2_ for SFRT. The depletion observations in different chemical environments showed a change in radiolysis and diffusion behavior, making it imperative to bring the enhanced depletion in the valleys and diffusion observations into the biological context to elaborate on the mechanisms for subsequent cell damage from this found radio-chemical contribution.

## VI. Acknowledgments

The authors would like to express their gratitude to the Division of Radiooncology/Radiobiology for providing the lab infrastructure and access to the Faxitron Multi-Rad225 Precision X-Ray machine. Furthermore, the authors thank Eric Arbes for the help with the dosimetry analysis. Additionally, C. Stengl acknowledges the funding from the Helmholtz graduate school for cancer research of the DKFZ, within their PhD program. The authors acknowledge funding by the Deutsche Krebshilfe (German Cancer Aid) Grant with the grant reference number 70115332.

## VIII. CRediT authorship contribution statement

**Constantin Schorling**: Conceptualization, Methodology, Formal analysis, Investigation, Data curation, Visualization, Writing - original draft, Writing - review & editing **Evelyn Rauth**: Conceptualization, Methodology, Investigation, Writing - original draft, Writing - review & editing. **Christina Stengl**: Conceptualization, Methodology, Writing - original draft, Writing - review & editing, Supervision. **Joao Seco**: Conceptualization, Writing - review & editing, Supervision.

