## Supplementary Material for "Experimental investigation of oxygen diffusion in the peak and valley region of minibeam patterns during X-Ray irradiation"

.

### Supplementary Section S.I. Dosimetry

With RW3 plates, a Semiflex Ionization Chamber 0.125 $cm^3$ Type 31010 (PTW, Germany) was positioned at the water equivalent height of the films under the phantom. Performing four sets of five absolute charge measurements for 2 min, the broad beam dose rate was established to be (6.12 $\pm$ 0.04) Gy/min. By irradiating films with a homogeneous dose (broad beam) at the same position for (0, 0.5, 1, 2, 5, 7, 10, 12, 15, 20) min, the optical density of the films was mapped to the corresponding dose by a 4th order polynomial fit, yielding a calibration curve. Further, three sets of minibeam irradiation for (0, 1, 5, 10, 15, 20) min with each collimator configuration were measured. Using the dose rate and the calibration curve, the dose in the peak and in the valley was calculated.

### Supplementary Section S.II. Minibeam Dose Pattern

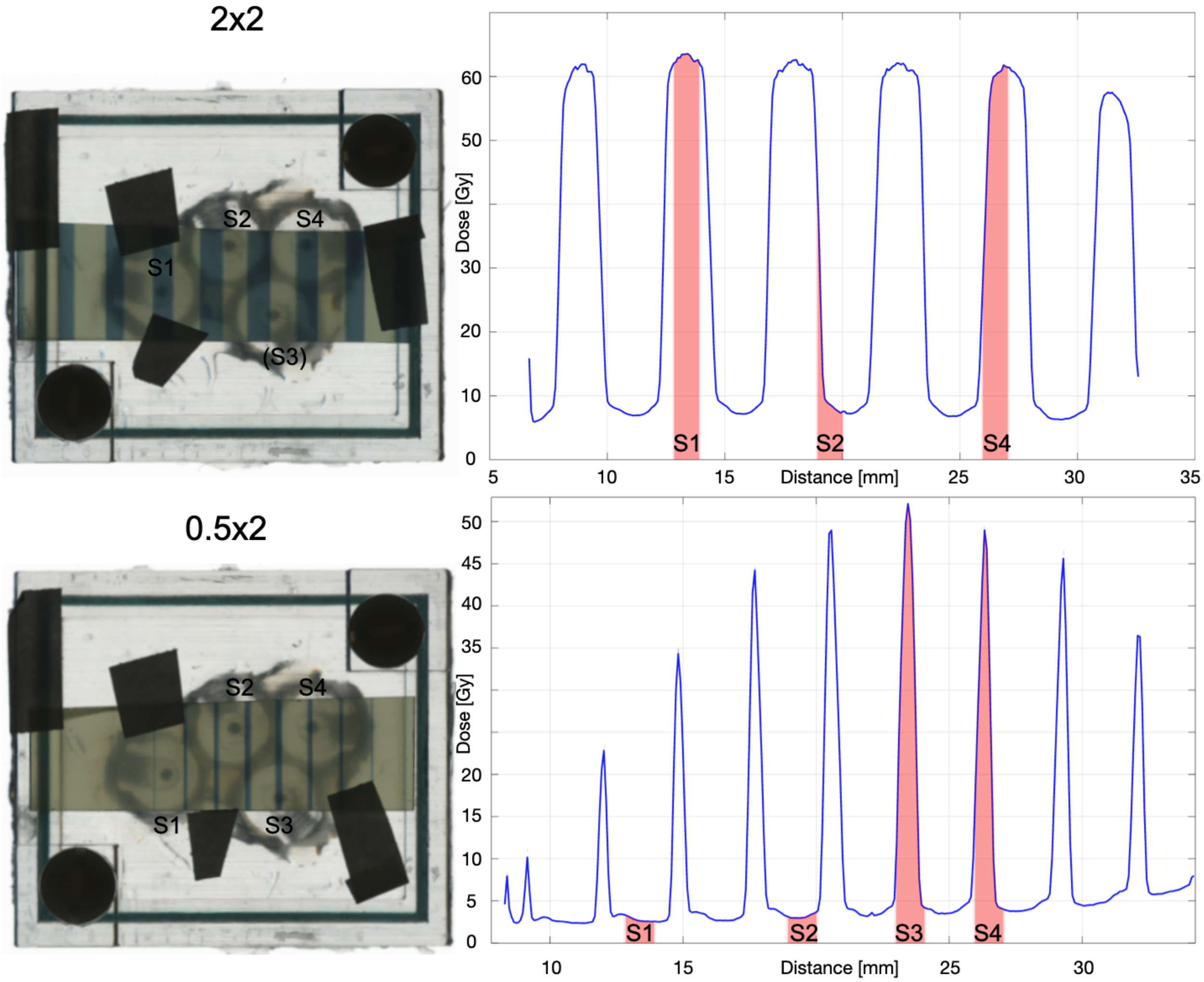


Supplementary Figure S1: Obtaining the minibeam dose pattern. Left: The scanned phantom with the labeled sensors and a film on top of the phantom to verfiy the sensor positions in the minibeam pattern. Right: Resulting dose pattern including the positioning of the sensors after film analysis. Top: Procedure for the 2.0 mm x 2.0 mm collimator configuration. Sensor S3 was excluded because it was not perfectly positioned in either a peak or a valley. Bottom: Procedure for the 0.5 mm x 2.0 mm collimator configuration.

### Supplementary Section S.III. $CO_2$ reactions

When a carbonated solution, i.e. water containing carbon dioxide or carbonic acid, is irradiated, carbon dioxide will also undergo radiolysis by being split into CO and $O_2$[1,2]. Furthermore, the by-products of the water radiolysis (e.g. $e^-_{aq}$, $HO^•$) will react with $CO_2$ and its by-products of dissolving carbonic acid[3]. Among others, the following reactions including their reaction rate constants $k$ are possible[3–5]:

$$HO^• + CO_3^{2-} \longrightarrow CO_3^{•-} + OH^-; \qquad k = 3.9 \times 10^8\ M^{-1}s^{-1} \tag{1}$$

$$HO^• + HCO_3^- \longrightarrow CO_3^{•-} + H_2O; \qquad k = 8.5 \times 10^6\ M^{-1}s^{-1} \tag{2}$$

$$HO^• + CO_2 \longrightarrow CO_3^{•-}; \qquad k < 1.0 \times 10^6\ M^{-1}s^{-1} \tag{3}$$

$$HO^• + CO \longrightarrow {}^•CO_2H; \qquad k = (1.2 - 2.0) \times 10^9\ M^{-1}s^{-1} \tag{4}$$

$$e^-_{aq} + CO_2 \longrightarrow CO_2^{•-}; \qquad k = 7.7 \times 10^9\ M^{-1}s^{-1} \tag{5}$$

$$e^-_{aq} + CO_3^{2-} \longrightarrow CO_2^{•-} + 2\,OH^-; \qquad k = 3.9 \times 10^5\ M^{-1}s^{-1} \tag{6}$$

$$e^-_{aq} + CO_2^{•-} \longrightarrow HCOO^- + OH^-; \qquad k = 9.0 \times 10^8\ M^{-1}s^{-1}. \tag{7}$$


### List of Figures


S1 SetUp . . . . . . . . . . . . . . . . . . . . . . . . . . . . . . . 2
